# Spatially modulated alpha-band activity does not mediate tactile remapping and fast overt orienting behavior

**DOI:** 10.1101/576850

**Authors:** José P. Ossandón, Peter König, Tobias Heed

**Affiliations:** Biological Psychology and Neuropsychology, University of Hamburg, 20146 Hamburg, Germany; Institute of Cognitive Science, University of Osnabrück, 49069 Osnabrück, Germany; Department of Neurophysiology and Pathophysiology, Center of Experimental Medicine, University Medical Center Hamburg-Eppendorf, 20251 Hamburg, Germany; Biopsychology & Cognitive Neuroscience, Faculty of Psychology and Movement Science, Bielefeld University, 33615 Bielefeld, Germany; Center of Excellence Cognitive Interaction Technology, Bielefeld University, 33615 Bielefeld, Germany

**Author notes:** Corresponding author: José P. Ossandón.

## Abstract

Posterior oscillatory alpha-band activity is commonly associated with spatial-attentional orienting and prioritization across sensory modalities. It has also been suggested to mediate the automatic transformation of tactile stimuli from a skin-based, somatotopic reference frame into an external one. Previous research has not convincingly separated these two possible roles of alpha-band activity. In particular, the use of delay paradigms, implemented to allow temporal evolution of segregable oscillatory brain responses to stimulus, motor planning, and response, have prohibited strong conclusions about a causal role of oscillatory activity in tactile-spatial transformations. Here, we assessed alpha-band modulation with massive univariate deconvolution, an analysis approach that disentangles brain signals overlapping in time and space. Thirty-one participants performed a delay-free, visual serial-search task in which saccade behavior was unrestricted. A tactile cue to uncrossed or crossed hands was either informative or uninformative about visual target location. Alpha-band suppression following tactile stimulation was lateralized relative to the stimulated hand over centro-parietal sensors, but relative to its external location over parieto-occipital sensors. Alpha-band suppression reflected external touch location only after informative cues, challenging the proposition that posterior alpha-band lateralization indexes automatic tactile transformation. Moreover, alpha-band suppression occurred ~200 ms later than externally directed saccade responses after tactile stimulation. These findings suggest that alpha-band activity does not play a causal role in tactile-spatial transformation but, instead, reflects delayed, supramodal processes of attentional re-orienting.

## Introduction

Oscillatory alpha-band activity exhibits modulation when human participants either expect or receive visual, auditory or tactile stimulation (Schürmann and Başar, 2001; Foxe and Snyder, 2011), and when they plan movements towards such stimuli (Buchholz et al., 2013). Alpha-band activity is usually suppressed contralateral to the attended side of space or contralateral to stimulation but can also be enhanced ipsilaterally. Both types of modulation result in hemispheric lateralization. Because alpha-band suppression is common across modalities and tasks, it has been suggested to reflect a supramodal spatial control mechanism (Klimesch et al., 2007; Jensen and Mazaheri, 2010).

However, for touch, it is not immediately clear how preparing for, or processing, a tactile stimulus should translate to alpha lateralization. The native reference frame of touch is skin-based, or somatotopic, that is, based on the body’s anatomy. Somatotopic coding differs from visual or, more generally, external coding because body parts frequently change position. Touch location in external space depends on limb posture: for instance, when the right hand crosses the midline, it is located in left external space. Indeed, alpha-band lateralization can reflect both somatotopic and external coding. Over somatosensory areas, alpha-band suppression depends on which hand is attended or stimulated, independent of posture (Buchholz et al., 2011, 2013; Schubert et al., 2015, 2018). In contrast, over occipito-parietal areas, alpha-band suppression depends on posture, suggestive of coding in external coordinates (Schubert et al., 2015, 2018).

Because occipito-parietal alpha-band modulation depends on the external location of touch, it could reflect the proposed supramodal spatial attention mechanism (Foxe and Snyder, 2011). Yet, because alpha-band suppression occurs in relation to somatotopic and external tactile location, it may, alternatively, be involved in tactile remapping, that is, the transformation of tactile-spatial information from somatotopic to external coding (Buchholz et al., 2011; Ruzzoli and Soto-Faraco, 2014; Schubert et al., 2018).

Previous research has not clearly dissociated these two possible roles of alpha-band activity, mainly because of two issues. First, tactile stimulation, and usually explicitly its spatial location, have been task-relevant in previous experiments. Therefore, alpha lateralization in those studies may have been due to attentional (re-)orienting and prioritization. Notably, external-spatial effects of touch on behavior have been observed even when a tactile stimulus is task irrelevant (Azañón et al., 2010; Ossandón et al., 2015), suggesting that tactile remapping occurs automatically. Thus, if alpha-band activity truly mediates tactile remapping, it should accompany the processing of task-irrelevant tactile stimuli, even if no observable behavior results.

Second, if alpha-band modulation played a causal role in tactile remapping, it should precede any externally oriented behavior in response to touch. Yet, whereas estimates for the time requirements of tactile remapping range from 150-300 ms (Yamamoto and Kitazawa, 2001; Azañón and Soto-Faraco, 2008; Overvliet et al., 2011; Brandes and Heed, 2015) alpha-band modulation has been demonstrated in tasks that implement long delays of >1s between attentional cue and tactile stimulus, or between tactile stimulus and response. During these delays, alpha-band suppression often develops gradually and is strongest at the end (Buchholz et al., 2011, 2013; van Ede et al., 2011, 2014; Bauer et al., 2012; Schubert et al., 2015). This time course may imply that alpha-band modulation reflects processes that result from, rather than cause, tactile-spatial transformation.

To scrutinize the role of alpha-band activity in tactile remapping and tactually induced attention, participants performed an overt visual search task in which the external location of a tactile cue was either informative or uninformative about visual target location, thus making the touch either task-relevant or irrelevant. Whereas task-irrelevant touch can nonetheless bias globally free-viewing behavior (Ossandón et al., 2015), the optimal strategy in the present experiment would be to follow informative and ignore uninformative tactile cues. Moreover, participants freely directed their gaze. By dissociating tactile, visual, and eye-motoric EEG activity using a massive univariate deconvolution approach (Ehinger and Dimigen, 2018), we investigated the dynamics of tactile processing as they presumably occur in real life, without artificial delays.

## Methods

### Participants

Thirty-six participants took part in the study after giving written consent. We excluded three of them from the analysis because they did not complete the experiment, and two because data were corrupted. We thus report on 31 participants (28 females; mean age: 25 years; range: 18 – 44; SD: 6.8). All experimental procedures were approved by the ethics committee of the German Psychological Society (TH 122014).

### Visual Stimuli

Figures 1A,B illustrate a visual search trial. The search stimulus was presented on a grey background that extended 30.3° horizontally and 24.1° vertically. It consisted of a grid of 6×8 white symbols with a single circle target located among 47 circle distractors for which a short, vertical line intersected the lower pole. This type of stimulus is known to require serial searching, as inferred from profiles of reaction time (Treisman and Souther, 1985; Zelinsky and Sheinberg, 1997). Stimuli were generated and presented with Psychtoolbox 3 (Brainard, 1997; Kleiner et al., 2007), executed on MATLAB R2007b (The MathWorks, Inc., Natick, Massachusetts, USA).

**Figure 1:**
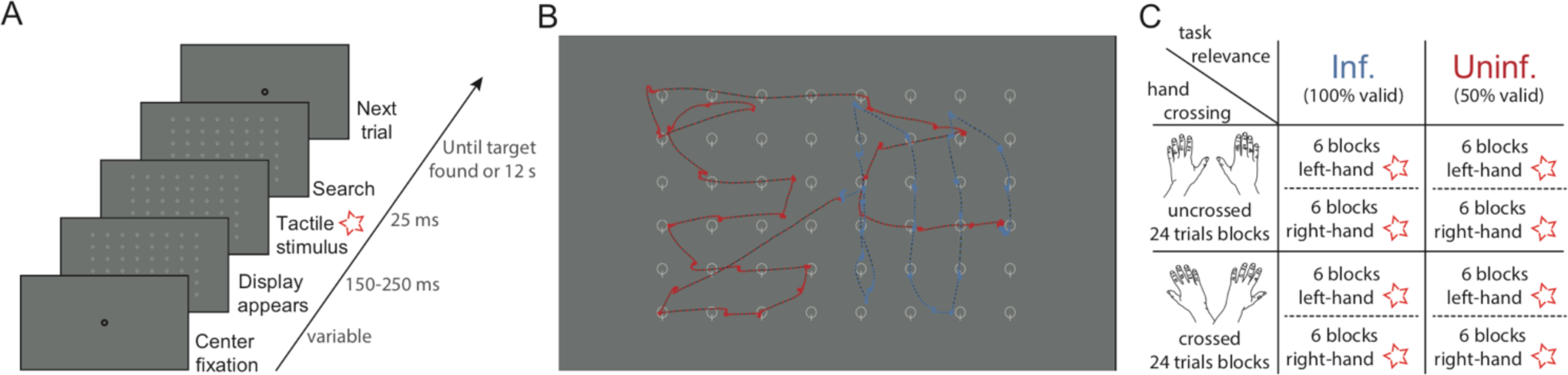
Experiment procedure. **(A)** Progression of a single trial. **(B)** Example of visual search. Stimuli consisted of a grid of 6×8 symbols with a single circle target (here in 4th row, 8th column) located among 47 distractors. Searches started at the center. Blue and red traces show eye-tracking data (each dot represents a sample acquired at 500 Hz) of one informative and one uninformative trial. In these two examples, the tactile stimulation was provided to the hand located below the right side of the screen. **(C)** Experimental conditions follow a 2 (left/right) × 2 (crossed/uncrossed) × 2 (informative/uninformative) fully factorial design. The star is used throughout the article as abbreviation for tactile stimulation. Informative and uninformative conditions were conducted as separate parts of the experiment. All other factors varied on a trial-by-trial basis.

### Tactile Stimulation

Tactile stimuli were 25 ms long, supra-threshold, 200 Hz vibrations, delivered to the back of the participant’s hand. Stimulation was produced by electromagnetic solenoid-type tactors (Dancer Design, St. Helens, UK) that were attached to the skin with medical tape. Tactors were driven linearly by the voltage generated via a Mini Piezo-Tactile Controller/Amplifier (Dancer Design, St. Helens, UK), which was directly controlled by the experimental computer’s audio and parallel-port outputs. To mask any noise that could originate from the stimulators, participants wore earplugs and white noise was played from a central loudspeaker located below the screen.

### Procedure

Fig. 1A illustrates the sequence of a single trial. Participants rested their hands comfortably on the table in front of them, with one hand below each side of the screen. Across blocks, hand position was altered between uncrossed and crossed postures, so that the right hand rested either under the right (uncrossed) or left side (crossed) of the screen, and vice versa for the left hand. Tactile cues occurred at the beginning of every trial, between 150 and 250 ms (uniform random distribution) after the appearance of the search display. The experiment was divided in two parts. In one part, the tactile cue was informative; participants were instructed that the search target was located on that side of the search display under which the stimulated hand was located. This tactile cue was 100% valid. This instruction was formulated externally, in that it pertained to the hands’ location in space, and not to the stimulated hand body side. In the second part, the tactile cue was uninformative; participants were instructed that tactile stimulation was unrelated to the search task. This tactile cue was, therefore, 50% valid. The order of the uninformative and informative experiment parts was balanced across participants.

Trials started with the appearance of the search display after the experimenter had confirmed that participants fixated a central fixation dot. The trial ended after the participant had fixated the target, operationalized as a cell of size 3.74° x 3.05° centered on the target, for more than 500 ms. Trials were aborted if no such target fixation occurred within 12s.

Fig. 1C illustrates the experimental conditions. Participants performed 48 blocks with 24 trials each. We, thus, collected 144 trials per combination of our three experimental factors Hand Crossing, Task Relevance, and Anatomical Side of Touch. Prior to testing, participants received a practice block of 16 trials without tactile stimulation.

### Eye-tracking

Eye movements were recorded with a video-based infrared Eyelink 1000 system (SR Research Ltd., Mississauga, ON, Canada), using the monocular remote tracking mode at 500 Hz sampling rate. Eye movements were defined using the systems’ default parameters, and the system standard 13-point calibration procedure was performed to achieve an average calibration error < 0.5° and a maximal calibration error < 1.0°.

### EEG acquisition

Electrophysiological data were recorded using Ag/AgCl electrodes with a BrainAmp DC amplifier (Brain Products GmbH, Gilching, Germany) with a sampling rate of 1000 Hz. 73 electrodes were placed according to the 10-10 system (Acharya et al., 2016), with location AFz serving as ground and the right earlobe as reference. Three electro-oculogram channels were placed in a triangular montage, with channels on the forehead and on the left and right infraorbital rim (Plöchl et al., 2012). The impedances of all electrodes were below 5 kOhm.

### EEG preprocessing

Data were preprocessed and analysed with MATLAB R2015a, using custom scripts and the third-party toolbooxes EEGLAB (Delorme and Makeig, 2004), Fieldtrip (Oostenveld et al., 2011), and Unfold (Ehinger and Dimigen, 2018).

The complete continuous EEG data was evaluated for the presence of artifactual segments and bad channels with an automatic, custom procedure based on amplitude, high-frequency noise and linear-trend thresholds (detailed description and analysis scripts available at https://osf.io/d7xc6). Channels or segments identified as artifactual were removed in subsequent analyses. We removed channels entirely when more than 15% of experimental data would have had to be discarded due to artifacts. This procedure excluded, on average, 0.77 channels per subject (16/9/4/1/1 participants had 0/1/2/3/4 channels removed, respectively). Excluded channels were replaced by interpolated channels based on spherical interpolation as implemented in EEGLAB’s *pop_interp* function. Next, we identified ocular and muscular artifactual components by independent component analysis and removed these components from the raw data. We identified eye-movement related components via an automatic algorithm based on the variance ratio between fixation and saccade periods (Plöchl et al., 2012). We defined muscle-related components as components in which total power above 20 Hz was larger than power below 20 Hz. We removed, on average, 24.2 (SD:7.3) independent components per subject.

### Data analysis

#### Behaviour

We evaluated the overall effect of cueing on reaction times with a three-way repeated measures ANOVA, with factors Anatomical Side of Touch (left/right), Hand Crossing (uncrossed/crossed) and Task Relevance (informative/uninformative). Individual main and interaction effects, consecutive two-way ANOVAs, and post-hoc comparison were evaluated with a significance threshold of 0.05, Bonferroni-adjusted for all possible multiple comparisons per-test.

We evaluated the effect of Task Relevance on the probability to make a saccade that ended on the cued side of the search display separately for uncrossed and crossed hand trials, using mixed-effect logistic models. We counted saccades in 50 ms bins from the moment of stimulation until 600 ms after, resulting in 12 time bins and, accordingly, 12 separate models. These logistic models were defined, in Wilkinson’s notation, by p(cued side) ~ Task Relevance + (1|subject). We report results as the probabilities associated with each condition (1/1+exp(-(model))) tested against the probability of no effect (p = 0.5) at a significance threshold Bonferroni-adjusted for multiple comparisons (12 time bins x 2 levels of Task Relevance = 24 tests).

#### Deconvolution models

Whereas we accounted for the well-known, massive artefacts caused by saccades by independent component analysis during preprocessing, the analyzed signal still contained multiple, overlapping signals generated by the appearance of the search display, saccade programming, and the sensory consequences of each new fixation. Moreover, ocular movements have variable latency, direction and amplitude. To separate all of these aspects from the effects of tactile stimulus processing, we combined massive univariate, linear deconvolution with generalized additive modelling (Ehinger and Dimigen, 2018). In massive univariate EEG modelling, regression of scalp electrical activity is performed independently for each sample and electrode. This approach has gained track over the last decade as a way to analyze EEG data obtained from complex setups that combine multiple experimental factors (Groppe et al., 2011; Pernet et al., 2011; Smith and Kutas, 2015a). Linear deconvolution is maybe best known from functional magnetic resonance imaging analysis (Dale, 1999). It determines time-extended effects of discrete experimental events that occur with varying temporal overlap during the experiment. For instance, here, a saccade and appearance of the search display are discrete events, but their respective effects on EEG signals potentially extend for several hundreds of milliseconds and, thus, overlap in time. Deconvolution allows to dissociate these overlapping signals (Smith and Kutas, 2015b; Ehinger and Dimigen, 2018). Moreover, generalized additive modelling allows accounting for non-linear effects, implemented here by using spline functions. This approach enabled us to integrate continuous regressors that account for saccadic movement amplitudes in the horizontal and vertical dimension. Finally, spatio-temporal clustering of regression weights in second-level analysis, that is, across participants, permits a data-driven statistical evaluation without the need to specify spatial or temporal regions of interest (detailed below).

The present analysis focused on alpha-band activity, because this frequency range has been consistently associated with the processing of tactile information, both for anatomical and external spatial coding. We down-sampled EEG signals to 250 Hz. The complete EEG dataset was bandpass-filtered with a sync zero-phase FIR filter (6dB cutoff at 7.8 and 16.1 Hz, 2.25 transition bandwidth) and Hilbert-transformed. Signal power at each sampling point was normalized in dB to its ratio with the respective channel mean power during a baseline period ranging from −450 to 0 ms before trial start, that is, at least 150 m before tactile stimulation.

We performed two analyses to examine the effects of tactile stimulation during visual search on alpha-band activity. The first analysis focused on the topography of alpha-band modulation relative to stimulus location, and we refer to it as the “topography analysis” from hereon. For this analysis, we re-coded channel topography relative to the anatomical site of stimulation. We flipped EEG channels for trials in which the left hand had been stimulated with respect to left and right (Buchholz et al., 2011; Schubert et al., 2015). This procedure effectively codes EEG signals as if all stimuli had occurred at the right hand, allowing us to pool trials across Anatomical Side of Touch to increase statistical power. We then applied a deconvolution model with effect-coding on the data of each subject. The model accounted for effects of tactile stimulation, search display appearance, and saccade initiation. For each event, effects were evaluated in a temporal window of 500 ms before to 800 ms after the respective event’s occurrence. The model contained the following predictors:

- for the effect of tactile stimulation, we entered main effects of factors Hand Crossing and Task Relevance, as well as their interaction (y ~ 1 + Hand Crossing x Task Relevance)
- for search display appearance, we entered an intercept (y ~ 1) relative to the time of search display appearance.
- for activity related to ocular movement, we entered an intercept and 10 spline predictors for the continuous variables of horizontal and vertical movement components, relative to the start of the saccade. We used splines instead of simple linear regressors because the effects of some saccade parameters, such as movement amplitude, on visual event-related potentials have been shown to be markedly non-linear (Dandekar et al., 2012; Kaunitz et al., 2014; Ehinger and Dimigen, 2018). Movement was computed as the position difference between saccade end and start (y ~ 1 + splines(xdiff, 10) + splines(ydiff, 10)). We reversed the sign of the horizontal eye movement component for left-hand trials, to account for flipping their EEG topography.

The second analysis focused on hemispheric differences of alpha-band activity, and we refer to it as the “hemispheric difference analysis” from hereon. We used the difference in power between homologous channels over the two hemispheres as dependent variable. This strategy reduced the number of channels to 33 (76 channels minus 10 midline electrodes, divided by 2). Furthermore, we did not pool left and right stimulus locations as in the topography analysis. Rather, keeping left and right stimulation separate allowed to derive contra- and ipsilateral effects by comparing right minus left stimulation. Accordingly, we applied a deconvolution model equivalent to that of our first analysis, but included a predictor of anatomical stimulation side and the respective interactions (y ~ 1 + Hand Crossing x Task Relevance x Anatomical Side of Touch).

#### Connectivity analysis

We hypothesized that information is transferred differently between primary somatosensory and posterior parietal regions depending on hand posture. To investigate information transfer, we analyzed connectivity measures between channels located above the respective brain regions that were also modulated by our task. Modulation of the respective channels by anatomical and external spatial codes has been previously reported (Buchholz et al., 2011; Schubert et al., 2015), and those studies demonstrated the related brain sources to include primary somatosensory and posterior parietal regions. We calculated the imaginary part of complex coherency in alpha-band activity as a way to measure oscillatory coupling that is not explained by spurious volume conduction (Nolte et al., 2004). First, we re-coded channels, as described above for the first time-frequency analysis, by flipping the EEG topography of left-hand trials. Next, we defined regions of interest (ROIs) from central and posterior channels that showed the strongest and longest modulation by anatomical or external spatial coding in the deconvolution analysis (central contralateral: ‘C3’; central ipsilateral: ‘C4’; posterior contralateral: ‘P5-P7-PO7’; posterior ipsilateral:‘P6-P8-PO8’). We then applied a sliding Fourier transform over the data of each trial, using a single Hanning taper with variable temporal window size consisting of 3 cycles at 9–15 Hz and 15 ms step size. Next, we calculated the imaginary part of coherency between each combination of pair of channels across ROIs, separately for each combination of Hand Crossing and Task Relevance. Finally, we determined a single value per ROI pair as the average across all possible between-ROI channel pairings.

#### 2nd-level analysis

We assessed statistical significance of alpha-band modulation with a second-level group analysis. To this end, the beta values of each model predictor were tested against zero at each time sample and channel with t-tests. We applied a cluster-based permutation test (Maris and Oostenveld, 2007) to control the elevated family-wise error-rate due to testing multiple channels and sampling points. We clustered all samples that resulted in a t-value with an associated p-value < 0.05 based on proximity on the scalp or in time, and the sum of the corresponding t-values was calculated for each of the resulting clusters. We compared empirical cluster values with the distribution of a 2000-samples permutation that we constructed by taking the maximal cluster value of each permutation iteration. Permutation iterations were obtained by randomly flipping, subject-by-subject, the sign of a given predictor across all channels and time samples (Good, 2000). This is based on the rationale that, if the estimated effects were not consistently different from zero, changing the sign of their values would results in similar clusters than the one obtained from the actual estimate. In contrast, if a factor results in a consistent effect, randomly changing predictor signs (across participants) would remove its effect. For any given predictor, we considered the determined empirical clusters significant at a p < 0.05 if their summed t-value was smaller than 2.5^th^ percentile, or higher than the 97.5^th^ percentile of the permutation distribution. To further control for testing multiple predictors, we applied a Bonferroni correction, dividing the obtained p-values by the number of predictors of the model (contra-ipsi models: 6 predictors, alpha = 0.008; difference models: 8 predictors, alpha = 0.005; connectivity comparisons: 6 predictors, alpha = 0.008).

### Open data and code accessibility

The code and model results used to produce the figures below are available at https://osf.io/d7xc6. The original EEG and eye-tracking datasets are available upon request.

## Results

Participants performed an overt visual search task in which they had to find a target among 47 distractors (see Fig. 1). At the beginning of each trial, one hand received a brief vibrotactile stimulus. This stimulus was either task-relevant and informed about the side of the search display on which the target was located, or it was uninformative and, thus, task-irrelevant. We asked how informative and uninformative tactile cues affected visual search behavior and EEG alpha-band activity. The hands were placed underneath the search display in an uncrossed or a crossed posture, allowing to determine whether modulation of behavior and electrophysiological responses were coded anatomically – that is, based on which hand was stimulated – or externally – that is, based on the side of space on which the stimulated hand was placed.

### Performance

Fig. 2A illustrates participants’ overall task performance and reaction time distribution. Performance was almost at ceiling, with the search target being acquired in 98.3% of trials within the 12s time limit. Across participants, the average search time was 2511 (sd: 500) ms. Fig. 2B illustrates the effect of tactile stimulation on search time. A repeated-measures ANOVA with factors Anatomical Side of Touch (left vs. right hand), Hand Crossing (uncrossed vs. crossed) and Task Relevance (informative vs. uninformative), revealed a significant main effect of Task Relevance (*F*_(30,1)_ = 132.3, p < 0.001) and a significant three-way interaction (*F*_(30,1)_ = 8.8, p = 0.006). Post-hoc comparison showed that search was 1156 ms faster with informative than uninformative cues (*t*_(30)_= −11.5, p < 0.001, ES: 2.06), a reduction of search time by 37%.

**Figure 2:**
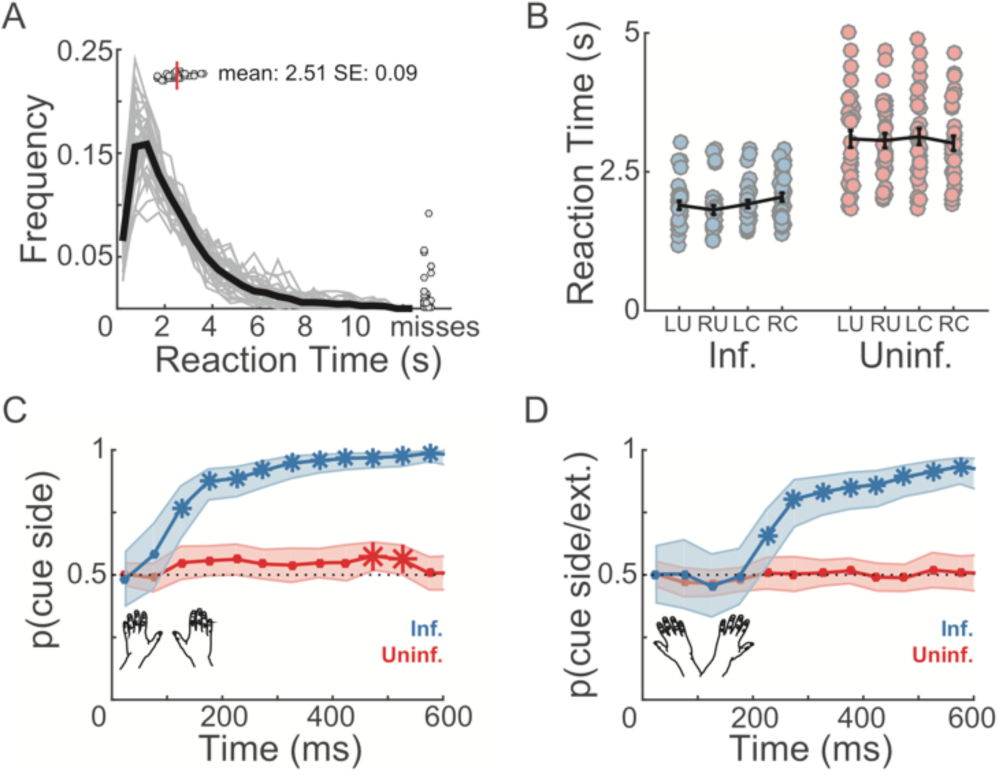
**(A)** Search performance. Thin gray lines show the search time distribution per participant; the black line reflects the group-average distribution. Gray circles on top indicate each participant’s mean search time; the red line indicates the average across participants. Gray circles on the right indicate the proportion of misses per participant. **(B)** Search time split by task relevance. Participants’ individual means are shown as gray circles. Error bars are condition mean +- s.e.m. **(C)** Probability to fixate the cued side in uncrossed-hands trials, binned into 50-ms segments following tactile stimulation. Blue and red lines indicate fixation probability for informative and uninformative trials, respectively. Shaded area reflects s.e.m. Asterisks demark probability different from 0.5 at a multiple-comparison Bonferroni-adjusted alpha for 24 tests (2 conditions × 12 bins of 50 ms width, ranging from 0-600 ms, see Methods). **(D)** As C, but for crossed-hands trials. Fixation side is coded relative to the stimulated hand’s location in space. Accordingly, the complement of the depicted probability corresponds to the probability of fixating on the side of the search display that corresponded with the anatomical body side of tactile stimulation.

We followed up on the three-way interaction with separate two-way ANOVAs for informative and uninformative cues. For informative tactile stimulation, there was a significant main effect of Hand Crossing (*F*_(30,1)_ = 10.7, p = 0.002) and an interaction of Hand Crossing and Anatomical Side of Touch (*F*_(30,1)_ = 14.3, p < 0.001). Post-hoc comparison indicated that hand crossing slowed search by 126 ms (uncrossed: 1856 ms; crossed: 1983 ms, *t*_(30)_= −3.2, p = 0.002, ES: 0.58). Furthermore, the effect of Anatomical Side of Touch depended on Hand Crossing: with uncrossed hands, search time was 84 ms faster after right as compared to left hand stimulation, though this difference was not significant after Bonferroni correction (*t*_(30)_= 2.22, p = 0.03, ES:0.4; corrected alpha significance level: 0.008). In contrast, with crossed hands, search time was significantly faster by 126 ms after left as compared to right hand stimulation (*t*_(30)_= 2.8 p = 0.008; ES: 0.5). This reversed hand effect suggests the observed processing advantage depended on the external location of touch in space, with that tactile stimulation on the right side of space being more effective than those on the left. In sum, touch was an effective spatial cue, and hand crossing resulted in a small but significant cost.

### Exploration patterns after tactile stimulation are biased only when stimulation is task-relevant

The effect of tactile stimulation on search time was paralleled by modulation of visual exploration behavior. Fig. 3A,B illustrates fixation positions of all participants for all experimental conditions. Fixations were centered on the search elements. In uninformative trials, exploration was distributed uniformly across items. In contrast, in informative trials, eye-movements were directed almost exclusively to the side of the screen indicated by the tactile cue in external coordinates. The only exceptions were a few saccades that were initiated before tactile stimulation.

**Figure 3:**
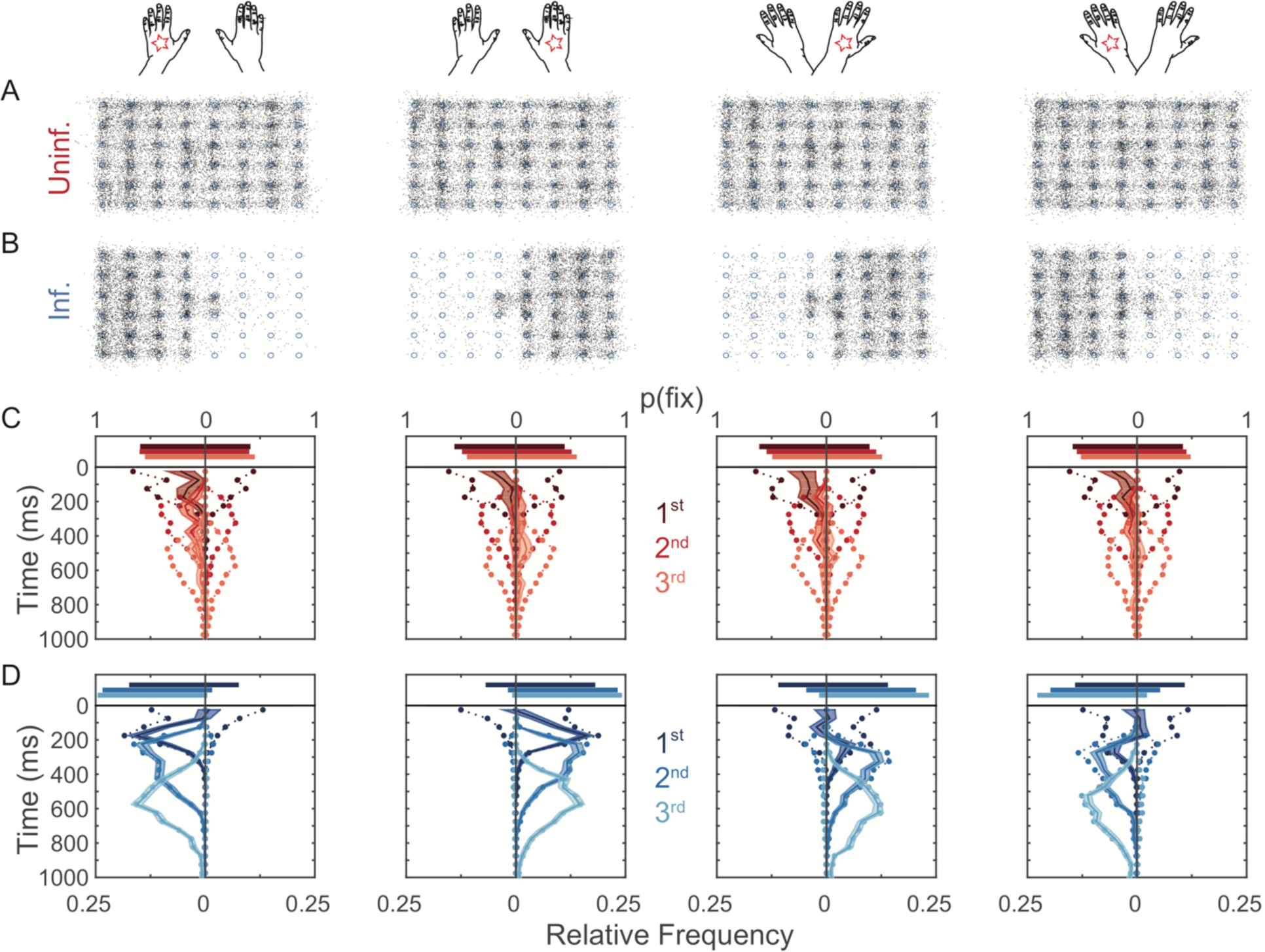
Exploration patterns and latency to move after stimulation. **(A,B)** Spatial fixation distribution of fixations for uninformative **(A)** and informative **(B)** cues, separately for each combination of Anatomical Side of Stimulation and Hand (columns 1-4). Panels display random subsamples of 50% of all participants’ pooled data to retain visibility of individual fixations. **(C,D)** Fixation probability and latency of the first three saccades according to the side of the screen on which they end, for uninformative **(C)** and informative **(D)** cues, separately for each combination of Anatomical Side of Stimulation and Hand. Bars in the upper section of each panel show the probability to fixate left or right for the first (dark), second (middle), and third (light) saccade after tactile stimulation. Dotted traces in the lower part of each panel illustrate the probability of saccades to end left or right over time, binned in 50 ms intervals, relative to the time of stimulation. Continuous lines with shaded area reflect the mean difference (+- s.e.m.,) between left and right relative frequencies.

This overall exploration pattern emerged directly following tactile stimulation. Fig. 3C,D, shows the probability to fixate and the latency distributions of the first three saccades after stimulation for informative and uninformative trials, grouped according to the hemifield in which the saccade ended. Saccade probability decreased from the moment of stimulation until around 100 ms after stimulation, to then increase again (see darkest traces in Fig. 3C,D); this pattern is typical for saccades following sensory stimulation and has been associated with saccadic inhibition due to stimulation (Ossandón et al., 2015). After this inhibitory phase, first saccades were biased to the left in uninformative trials, as it is usually the case during exploratory free-viewing (Ossandón et al., 2014, 2015). Second and third saccades following stimulation were balanced across hemifields. In informative trials, second and third saccades ended on the side of the display that had been cued by the tactile stimulus. The preference for saccades to the cued side of the search display (continuous lines in Fig. 3C,D) became apparent after ~100 ms with uncrossed hands, but not until ~200 ms with crossed hands.

We statistically tested the modulation of saccade end-points by tactile stimulation with mixed-effect logistic models for the averaged fixation probability of every 50-ms bin from the moment of tactile stimulation until 600 ms after stimulation, separately for uncrossed and crossed hand trials. For informative trials with uncrossed hands (Fig. 2C), the first time bin in which fixation probability to the tactually cued side differed statistically from chance was the 100-150 ms time bin. Fixation probability increased to above 95% in the following time bins, statistically confirming the above reported observations. In contrast, exploration was unbiased by the cue in uninformative trials, with the exception of the interval between 450-550 ms, in which fixation probability was slightly but significantly biased towards the side indicated by the uninformative cue.

For crossed hands (Fig. 2D), saccade probability towards the tactually cued side refers to instances in which tactile location had been transformed from anatomical to external location. For informative trials, the first time bin in which saccades were more probable towards the tactually cued side of the search display was the 200-250 ms time bin. Fixation probability increased to above 90% in the following time bins. Exploration was unaffected by the cue in uninformative trials. At no time did we observe a bias towards the anatomical side of stimulation in crossed hand trials. This finding suggests that saccades were guided by the external location of tactile cues.

### Anatomically and externally coded alpha-band EEG responses to tactile stimulation

Analysis up to this point demonstrated that tactile stimulation affected visual search behavior, and that this modulation depended on the relevance of tactile stimulation for the search task. The focus of the present paper, however, was on the modulation of alpha-band activity by touch and saccades in the context of visual exploration. Previous research has shown that both the expectation of tactile stimulation, as well as tactile stimulation proper correlate with decreases in oscillatory alpha-band activity that can be related to coding in different reference frames. Here, too, we observed that modulation of oscillatory activity by our task manipulations was distinct to the alpha-band range around 10 Hz (see Fig. 4A), evident both during baseline and after stimulation (see Fig. 4B).

**Figure 4:**
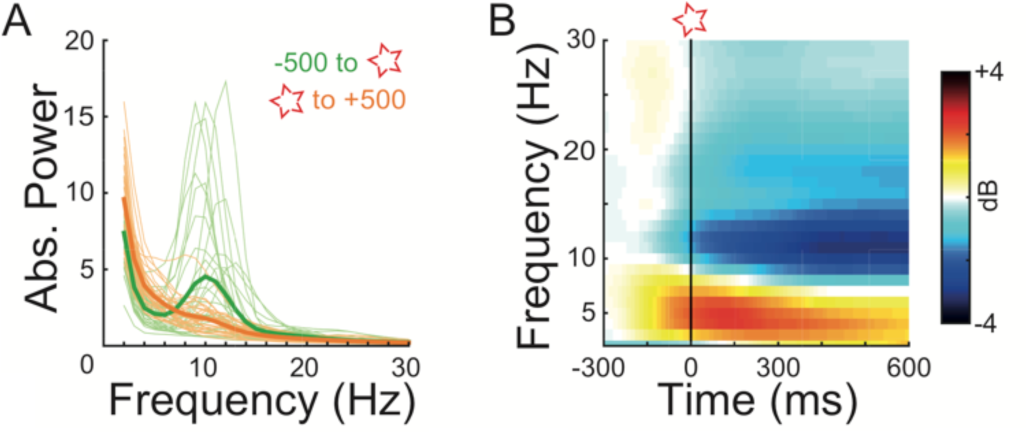
Absolute and relative spectra of all channels in all conditions. **(A)** Absolute power spectra of each subject (thin lines) and averages (thick lines) for the baseline period between - 500 to 0 ms prior tactile stimulation (green) and for the period between 0 to +500 ms after stimulation (orange). **(B)** Time-frequency chart aligned to the moment of tactile stimulation in dB with respect to a baseline period between −600 and −400 ms before stimulation (window size: 500 ms; padding: 1000 ms; single hanning taper; moving step: 15 ms).

We statistically evaluated the effect of tactile stimulation on alpha-band activity, restricted to 9-15 Hz, with a massive-univariate deconvolution modeling approach that used predictors for tactile stimulation, the start of the search display, and saccade programming on EEG activity. The deconvolution model isolates effects of tactile stimulation in the context of visual processing and unrestricted saccades across extended periods of time. As a critical feature, we modeled horizontal and vertical saccade displacement with a set of splines to remove any confounding effects of the systematic behavioral saccade direction biases that we had identified in search behavior.

We performed two analyses on alpha-band activity, one focusing on the topography of alpha-band modulation across the scalp, and the other focusing on hemispheric differences of alpha-band activity.

The topography analysis recoded all data channels as if all tactile stimuli had been applied to the right hand (see Methods). Cluster-based permutation testing identified significant clusters of power modulation for the predictors associated with the tactile stimulation model intercept, the main effects of Hand Crossing and Task Relevance, and of their interaction. Fig. 5A displays the alpha-band modulation associated with each of these predictors as topographic maps. The intercept was negative surrounding the time of stimulation in most channels both ipsi- and contralaterally (Fig. 5A, top row, significant cluster from −228 to +800 ms, p < 0.0005), indicating that tactile stimulation was accompanied by a general alpha-band power decrease. The fact that this decrease began prior to tactile stimulation was probably related to the predictability of tactile stimulation to occur in temporal vicinity of trial start, even if it was jittered. We observed a similar, global suppression effect for the factor Task Relevance that was stronger in informative than uninformative trials, but offset by ~400 ms (Fig. 5A, 2nd row, cluster from +392 to +800 ms, p < 0.0005). Hand Crossing was associated with alpha-band suppression in a lateralized cluster ipsilateral to the stimulated hand (Fig. 5A, 3rd row, cluster from +156 to +796 ms, p = 0.0025); that is, when the hands were crossed, alpha-band suppression was contralateral to the external location of the tactile stimulus. Finally, the interaction between the two factors was accompanied by another cluster ipsilateral to the stimulated hand (fig. 5A, 4th row, cluster from +324 to +800 ms, p = 0.005), indicating that the Hand Crossing effect was larger for informative than uninformative trials. The effect of Hand Crossing and its interaction with Task Relevance can be more easily appreciated in Fig. 5B,C, which show averaged alpha-band power for each factor combination, as well as the difference between uncrossed and crossed postures, separately for informative and uninformative trials. Contralateral to the anatomical side of tactile stimulation, alpha-band activity was suppressed both at central and posterior channels, independent of hand posture. However, posture did affect central and posterior suppression ipsilateral, with stronger suppression with crossed than with uncrossed hands. Note, that the anatomically ipsilateral hemisphere is, at the same time, contralateral to external stimulus location with crossed hands.

**Figure 5:**
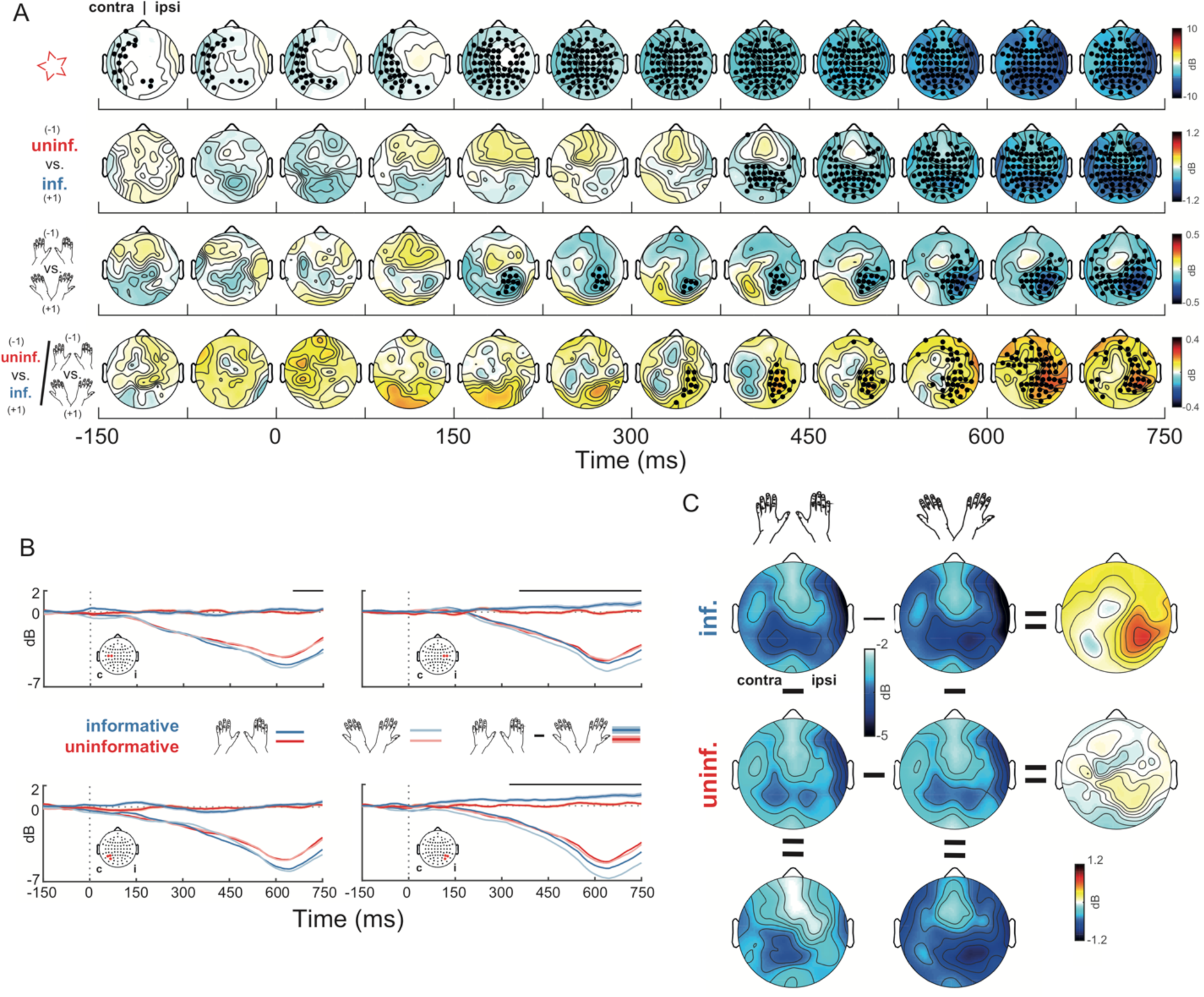
Modulation of alpha-band (9–15 Hz) power by tactile stimulation during visual search. **(A)** Average EEG topographies of the power assigned to relevant predictors of the deconvolution model in which all stimuli were coded as having occurred on the right hand, in time steps of 75 ms. Time 0 is the moment of tactile stimulation. From top to bottom, the rows show the tactile stimulation intercept, main effects of Task Relevance and Hand Crossing, and their interaction. Dots indicates sensors that have been assigned to a significant spatio-temporal cluster by cluster-based permutation testing with at least one sampling point. Gray dots indicate p < 0.05 corrected for the multiple electrode and time samples tested per factor; black dots indicate significance after correcting for the number of permutation-tests per model (see Methods). **(B)** Alpha-band suppression following tactile stimulation for central (top panels) and posterior (bottom) channels, separated by hand posture and task relevance. Lines with shaded areas depict the difference between uncrossed and crossed hands and, thus, reflect the interaction term in A. Horizontal lines on top of the plots show when any of the included channels (see insets) form part of a significant cluster. **(C)** Topographies of alpha-band suppression for the different experimental conditions and respective contrast in the interval between +300 and +750 ms after stimulation, in which the interaction between Hand Crossing and Task Relevance was significant.

Multiple studies have suggested that differences between alpha-band power between the two hemispheres is indicative of attentional lateralization (Foxe and Snyder, 2011). Therefore, we performed a second analysis to directly assess the disbalance of alpha-band activity between the two hemispheres. We statistically analyzed the difference of homologous channels of the left minus the right hemisphere, for instance C3 minus C4, with deconvolution modeling. In this analysis, we did not collapse across left and right hand stimulation, to allow assessing effects of contra- and ipsilateral stimulation as left minus right stimulation trials (see Methods). Accordingly, the model comprised the same predictors for Hand Crossing and Task Relevance as the topography analysis, but in addition included predictors for the main effect of Anatomical Side of Touch as well as the interactions with this factor. Fig. 6 illustrates the topographies for predictors that were significantly modulated by tactile stimulation. The main effect of Anatomical Side of Touch (Fig. 6, top row) was evident as stronger alpha suppression for a circumscribed cluster of central channels contralateral to the stimulated hand (cluster from +64 to +800 ms, p < 0.0005). Given that the dependent measure of the analysis was the difference of left minus right channels, this suppression effect indicates stronger suppression anatomically contralateral to tactile stimulation. There were no significant clusters for the interaction of Anatomical Side of Touch and Task Relevance, suggesting that anatomical coding was not modulated by task relevance. The interaction of Anatomical Side of Touch and Hand Crossing (Fig. 6, middle row) and the three-way interaction of Anatomical Side of Touch, Hand Crossing, and Task Relevance (Fig. 6, bottom row), revealed significant clusters of alpha suppression when corrected per predictor (Side x Crossing: +276 to +596 ms, p = 0.049, three-way interaction: +324 to +600 ms, p = 0.019). However, these clusters did not survive correction for multiple factors. We note, that they were spatially congruent with the clusters revealed by our first analysis. Thus, even if they give only weak additional support to those results, importantly they did not render any conflicting results.

**Figure 6:**
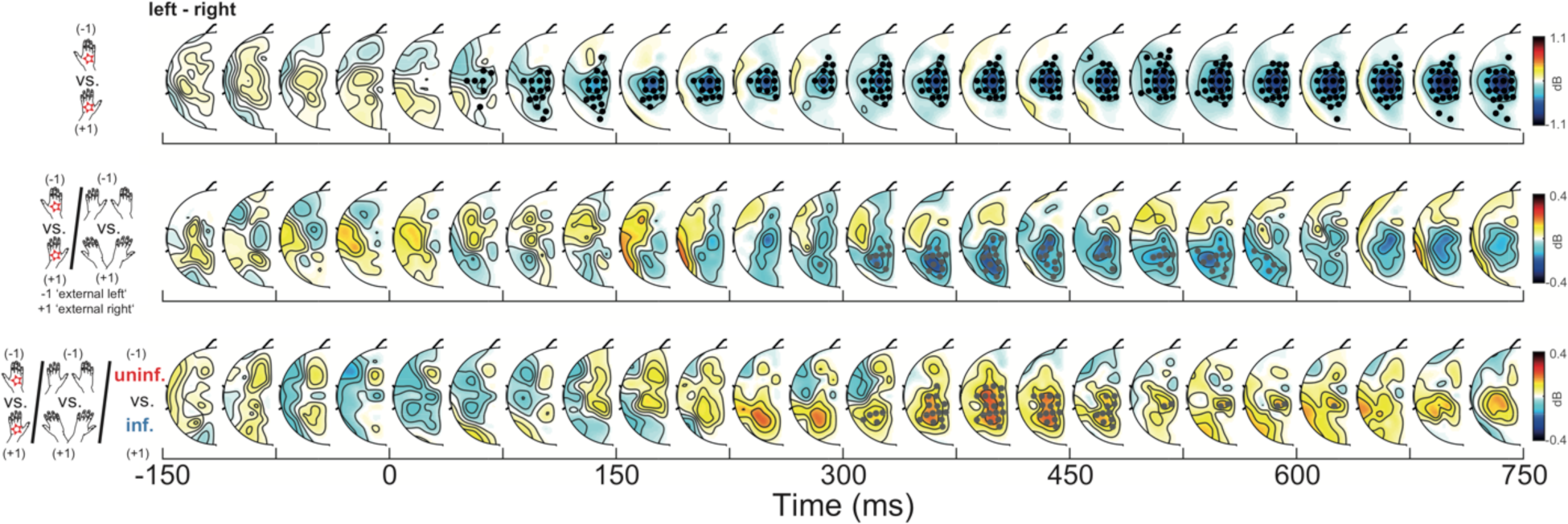
Modulation of lateralized alpha-band power by tactile stimulation during visual search. Alpha-band activity was subtracted (left minus right) between homologous channels. From top to bottom, the rows show the effect of anatomical-hand stimulated, the interaction of Anatomical Side of Stimulation and Hand Crossing, and the three-way interaction of Anatomical Side of Stimulation, Hand Crossing, and Task Relevance. Topographies show the left hemisphere, from which right hemisphere activity was subtracted. Midline electrodes were omitted. Each topography depicts the average of a 75 ms time interval. Time 0 is the moment of tactile stimulation.

In summary, the results of the two analyses revealed suppression of alpha-band activity following tactile stimulation. These suppression effects were independent of hand posture at contralateral central channels, indicating that alpha-band modulation was affected by anatomical coding. In contrast, alpha-band activity was modulated by hand posture at anatomically ipsilateral central and posterior channels, suggesting that touch affected neural activity in an external spatial code. This latter effect was evident only in informative trials, that is, when the stimulus location was task-relevant.

### Alpha-band EEG responses to the search display and saccades are discernible from responses to touch

Our deconvolution model accounted for visual and saccade effects to isolate effects of tactile stimulation. This separation of processing related to touch and eye movement is particularly relevant because visual, saccade, and tactile processing have all been associated with regions in posterior parietal cortex (Andersen and Cui, 2009). In fact, transcranial magnetic stimulation has targeted identical intraparietal regions to modulate alpha-band activity related to both visual and tactile processing (Romei et al., 2010; Ruzzoli and Soto-Faraco, 2014). The deconvolution model can tease the different processes apart because they occur at independent points in time and extend over stretches of several hundred milliseconds. We scrutinized whether parietal alpha-band activity was indeed assigned to our predictors of interest, as well as to other predictors for which parietal activity should be evident. Fig. 7A illustrates that alpha-band activity was suppressed by the appearance of the search display. Statistically significant clusters were present only in the topography analysis, in which alpha-band activity was tested over the two hemispheres. Alpha-band power was globally suppressed from −256 to +648 (p < 0.0005) first globally and then more locally to posterior and right side electrodes, and enhanced again from +416 to +796 ms (end of the analysis window, p < 0.0005), first in lateral electrodes in the left hemisphere and then globally. In the hemisphere difference analysis, clusters obtained with permutation tests did not survive correction for multiple testing.

**Figure 7:**
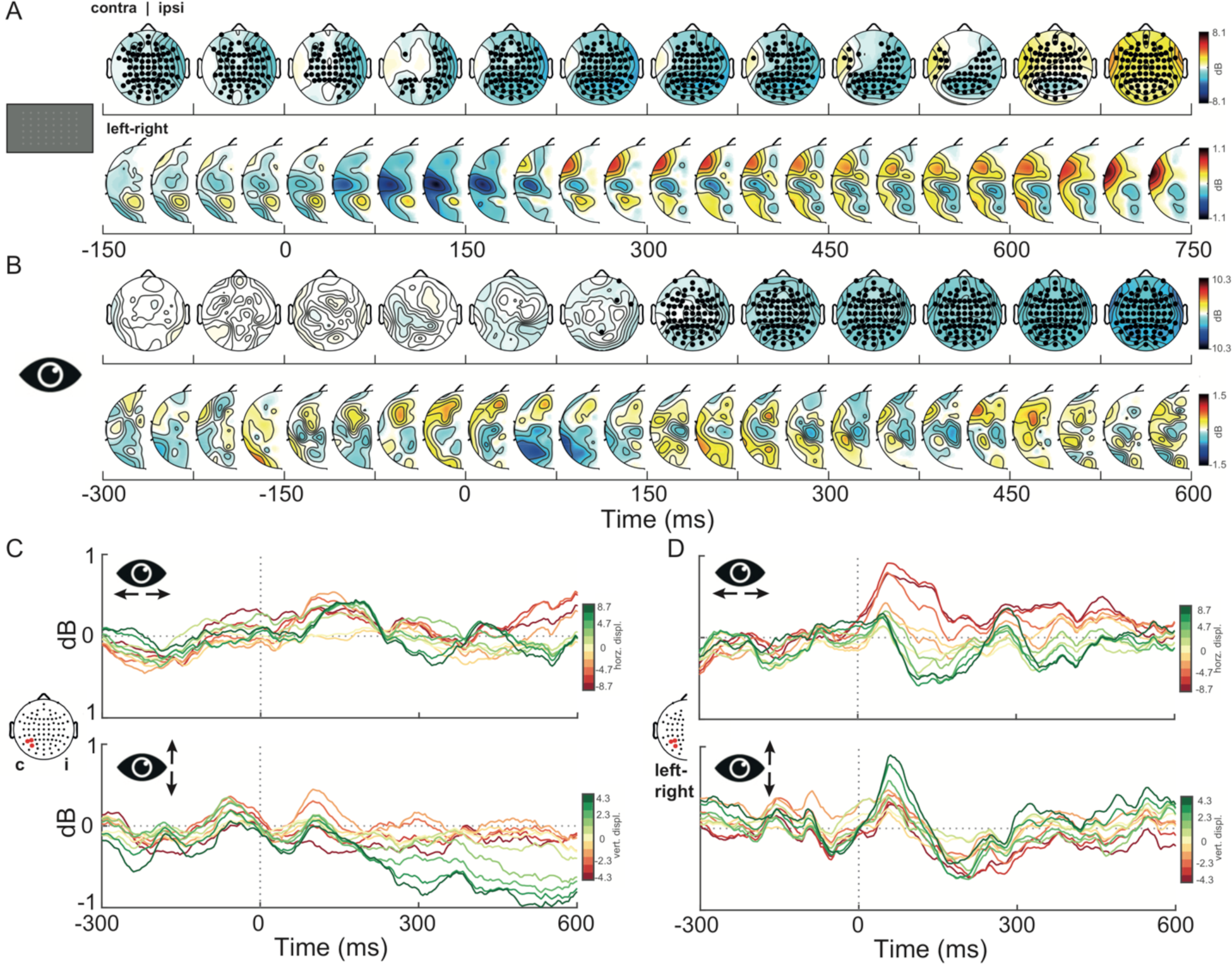
Alpha-band (9–15 Hz) modulation by the occurrence of the search display and saccades. **(A)** Effect of display appearance; upper row: first analysis (see Fig. 5 and main text); lower row: second analysis (see Fig. 6 and main text) **(B)** As A, but for the saccade event intercept. (C,D) Effects of differently sized vectors of horizontal (top panels) and vertical (bottom panels) saccadic displacement obtained from spline predictors in the deconvolution modelling for the topography **(C)** and hemisphere difference **(D)** analyses. Traces show activity of the summed spline predictors, averaged over those posterior channels that were significantly modulated.

Once the search display was presented, participants were allowed to saccade without restriction, resulting in the production of a sequence of fixations and saccades. Therefore, to disentangle the effects of tactile stimulation from the ones related to saccade programming and subsequent visual processing, we included all saccadic events that started within 800 ms of trial start. Fig. 7B shows the saccade event intercept for both contra-ipsilateral (top row) and interhemispheric difference (bottom row) analyses. Only the topography analysis revealed significant clusters of alpha-band activity modulation, with a global cluster of inhibition from +132 to +796 ms (end of the analysis window, p< 0.0005). These results imply that each new saccade event further suppressed alpha-band activity above and beyond any suppression due to the appearance of the search display as well as tactile stimulation. It is noteworthy that we did not find any alpha-band modulation prior to saccade initiation. This observation suggests that alpha-band activity does not reflect saccade programming, neither globally, nor lateralized, that is, in hemispheric activation differences. This lack of alpha-band modulation prior to saccades is in stark contrast with a modulation of alpha-band activity that depended on both horizontal and vertical distance of the performed saccade (see Fig. 7C,D). Thus, with each saccade, alpha-band activity was not just modulated globally, as evident by the significant saccade intercept predictor in our model (see Fig. 7B), but alpha-band activity was sensitive to the size of the specific saccade parameters. Alpha-band activity was suppressed in the hemisphere contralateral to saccade direction, with stronger suppression the larger the saccade. This graded effect was long-lasting, but was strongest about 50-200 ms after the saccade (see Fig. 7D). Notably, as for the other saccade predictors, no modulation of alpha-band activity was evident prior to saccade initiation. This suggests that the graded effect of saccade amplitude was a result of saccade execution, and not related to the planning of the saccade.

### EEG connectivity suggests that external coding is mediated by ipsilateral somatosensory cortex

Next, we asked how information is routed from somatosensory regions to posterior parietal cortex. Given the prominent effects in the alpha range, we hypothesized that externally coded alpha-band modulation may be mediated by oscillatory coupling with somatosensory cortex. More specifically, we conjectured that the somatosensory cortex contralateral to tactile stimulation, in which tactile information first arrives in cortex, would exhibit coupling with the posterior regions contralateral to the tactile stimulation’s external location. Accordingly, with uncrossed hands, coupling should be intra-hemispherical, between somatosensory and posterior parieto-occipital cortex of the same hemisphere. In contrast, with crossed hands, anatomical and external location belong to opposite hemifields, and therefore coupling should be evident cross-hemispherically, between the somatosensory cortex contralateral, and the posterior parieto-occipital cortex ipsilateral, to the anatomical side of tactile stimulation. Notably, connectivity and signal power reflect independent neuronal mechanisms. Therefore, it is possible that, even if alpha-band power rose late after tactile stimulation, alpha-band connectivity may raise significantly earlier, and indicate a causal role of this frequency band for tactile-spatial transformations through connectivity rather than local power modulation.

We tested connectivity between the specific central and parietal channels of the two hemispheres in which we had observed anatomically and externally coded spatial modulation of alpha-band activity, as evident in alpha-band modulation in response to our experimental factors of Hand Crossing and Task Relevance. We assessed connectivity as the imaginary part of the complex coherency. Before tactile stimulation, connectivity between central and parieto-occipital channels was enhanced intra-hemispherically, but not cross-hemispherically, independent of hand posture and task relevance of tactile stimulation (Fig. 8B,D). Cross-hemispheric coupling was, however, evident after tactile stimulation. Yet, contrary to our hypothesis, we did not observe cross-hemispheric coupling between central and posterior channels (Fig. 8C,D). Instead, cross-hemispheric coupling was evident between the two parieto-occipital regions, and only during informative trials (Fig. 8F, difference cluster from +400 to +800 ms, p<0.001). This cross-hemispheric coupling was accompanied by intra-hemispheric coupling of the ipsilateral somatosensory and parieto-occipital channels (Fig. 8E). Although ipsilateral coupling was present in informative and uninformative trials, a modulation of connectivity by hand posture was apparent only in informative trials.

**Figure 8:**
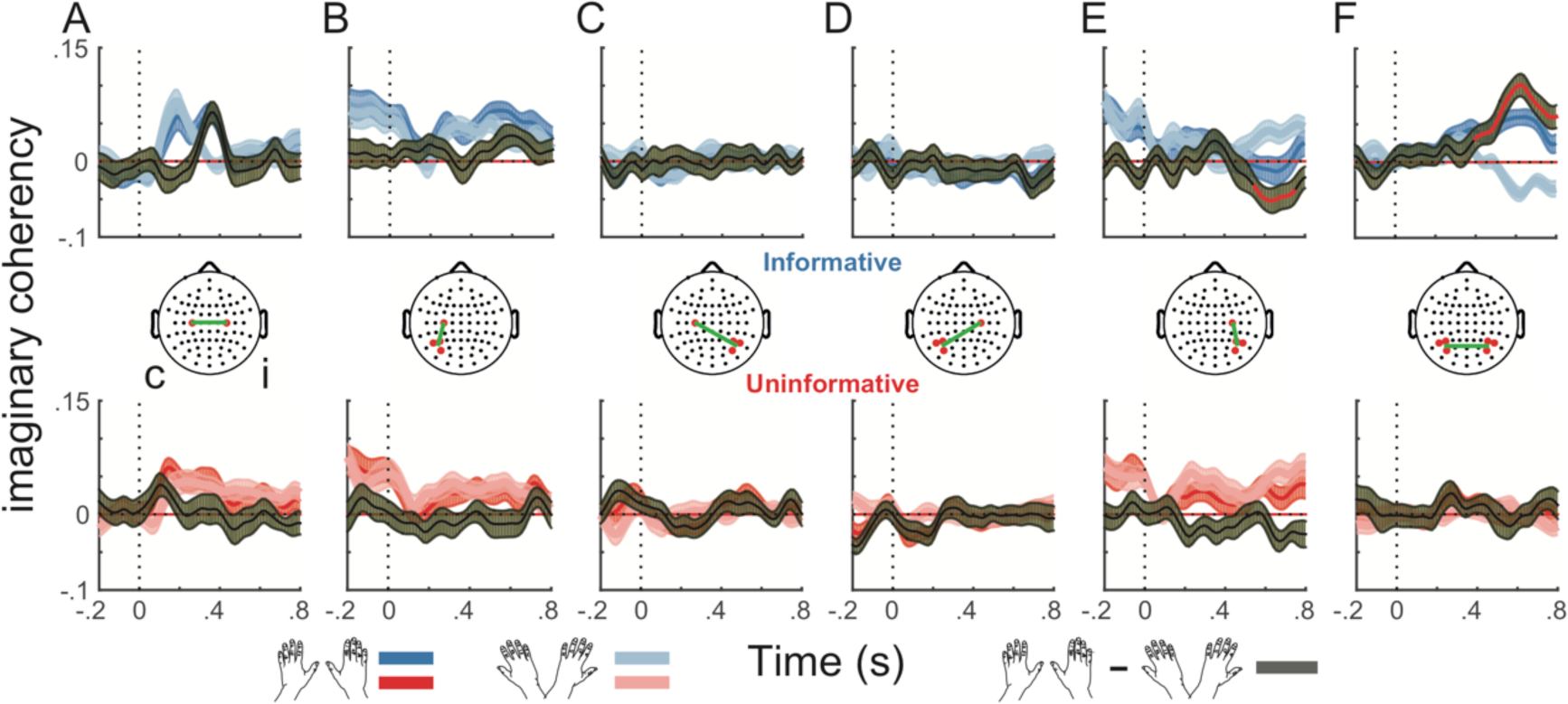
Alpha-range coupling between central and parietal regions, assessed as the imaginary part of coherency. Top and bottom plots show the results for informative and uninformative trials respectively. Different columns illustrate the coupling between each pair of regions of interest. Dark and light lines indicate uncrossed and crossed hand data, respectively. Black line shows the difference between postures, with a red line denoting a significant temporal cluster. Shaded areas represent s.e.m.

Together, these results are incompatible with direct transfer of information by alpha-band coupling between somatosensory and posterior parieto-occipital areas across hemispheres as a means of transforming anatomically coded into externally coded tactile information to guide attention. Instead, they suggest that interhemispheric transfer of tactile information is mediated in a first step by coupling between somatosensory areas, later followed by intrahemispheric coupling between somatosensory and posterior parieto-occipital areas.

## Discussion

The present study aimed at characterizing the role of alpha-band modulation during the processing of tactile-spatial information. In particular, we asked whether the occurrence and timing of alpha-band modulation supports the notion that this brain signal may be a mediator of tactile-spatial transformations. Our study revealed three main results: first, touch affected saccade behavior exclusively in an external reference frame, and only when it was task-relevant. This effect on behavior was evident within 100 ms for uncrossed hands, and within 200 ms for crossed hands. Second, alpha-band suppression was modulated somatotopically in contralateral somatosensory cortex, but externally in ipsilateral somatosensory and posterior parietal cortex. External-spatial effects of touch on alpha activity and interhemispheric connectivity between somatosensory and posterior parietal regions were evident only when touch provided task-relevant information. Third, saccadic behavior was modulated by touch before a change of alpha-band suppression was evident in the EEG signal. Together, these findings are inconsistent with the notion that alpha-band activity directly reflects tactile-spatial transformation and suggest that the involvement of this oscillatory signal, instead, reflects supramodal, attention-related consequences of remapped tactile information.

### Behavioral correlates of tactile stimulation

Tactile spatial information guided overt visual search when it was informative. This effect was fast and consistent: after 100 ms, saccades were biased towards the external side of touch; subsequent saccades remained almost exclusively on the cued side. When the hands were crossed, these effects were slower by another 100 ms, but nonetheless directed externally. This finding replicates our observation in a free-viewing task of natural scenes (Ossandón et al., 2015). While a purely external effect of touch on saccades is in agreement with previous studies (Groh and Sparks, 1996; Blanke and Grüsser, 2001; Overvliet et al., 2011; Buchholz et al., 2012), both saccades and reaches towards tactile locations can exhibit trajectories that initially deviate towards the anatomical side of the tactile event (Groh and Sparks, 1996; Overvliet et al., 2011; Brandes and Heed, 2015). Note that, in those tasks, the tactile stimulus location was the movement goal. The present and our earlier study (Ossandón et al., 2015) suggest, in contrast, that global biases during free-viewing, in which the tactile stimulus is not the goal of the subsequent movements, depend solely on an external reference frame. However, in contrast to our previous report in which tactile stimulation was task-irrelevant and the elicited bias was inconsequential, participants did not exhibit any bias following uninformative cues in the present study. This finding indicates that tactile spatial biasing is not mandatory and can be suppressed.

Previous estimates of when external tactile information first guides behavior have been inconsistent, ranging from about 150 to 360 ms (Yamamoto and Kitazawa, 2001; Azañón and Soto-Faraco, 2008; Overvliet et al., 2011; Brandes and Heed, 2015). At 100 ms response time, externally directed saccade responses were considerably faster here, and conflict resulting from crossed limbs was resolved after 200 ms, consistent with our previous report about hand reach corrections to tactile stimuli (Brandes and Heed, 2015). The time from programming a saccade to its execution has been estimated to lie around 100 ms, based on countermanding and saccade inhibition tasks (Hanes and Carpenter, 1999; Reingold and Stampe, 1999). Likewise, we and others have reported tactile saccadic inhibition effects within 100 ms from stimulation (Åkerfelt et al., 2006; Ossandón et al., 2015). Prior to 100 ms, saccades are directed randomly to all locations on the screen; afterwards, saccade direction integrates tactile stimulus location. This suggests that saccades cannot be modified within the last 100 ms before they are initiated, and thus the specification of the saccade’s spatial goal must have taken place before that time. Strictly speaking, then, saccades following touch to uncrossed hands in our study suggests almost immediate transformation of somatotopic into external touch location. However, one must bear in mind that our time estimates are averages across many trials, and that the stimulation only affected some saccades at 100 ms, becoming more consistent only at later time points. In a previous study, we compared when participants initiated a turn of a straight hand reach towards a visual or a tactile stimulus, presented in-flight (Brandes and Heed, 2015). Tactually evoked turns were only 20 ms slower after tactile stimulation to uncrossed limbs than to visual stimuli at identical locations, thus suggesting a very short estimate for the computation of an external-spatial location of touch. The present results further support this notion.

### Lateralized alpha-band suppression in response to tactile stimulation

Previous work has consistently shown that spatial cueing of tactile stimulation is followed by central and posterior parietal alpha-band suppression in the interval between the cue and presentation of the tactile stimulus (Jones et al., 2010; Haegens et al., 2011; van Ede et al., 2011; Bauer et al., 2012). Comparison of hand postures revealed that these alpha-band suppression effects depend on the cued hand over central electrodes, but that they additionally reflect the cued space at posterior parietal electrodes (Schubert et al., 2015).

Other studies have, instead, investigated alpha-band modulation after presentation of a tactile stimulus. In one study, participants received tactile stimulation on the index or little finger of one hand while fixating the same hand’s middle finger (Buchholz et al., 2011). While planning a saccade to the tactile location, posterior parietal alpha-band activity was suppressed in the hemisphere opposite to finger location relative to gaze, as it has been reported for visual paradigms as well (Gutteling et al., 2015). In another study, touch to uncrossed and crossed hands also resulted in externally coded parietal alpha-band suppression (Schubert et al., 2018).

Here, a tactile cue was used to modulate attentional and motor processing of an ongoing task that involved frequent saccades. As such, our study cannot be directly compared to previous experimental paradigms that have either asked participants to prepare for a touch or investigated tactile processing that elicited a single response towards a tactile location. Nevertheless, alpha-band suppression was similarly lateralized as in tasks that require attention in preparation for tactile stimulation. Crucially, parieto-occipital lateralization in external coordinates occurred only when touch was informative. Given the absence of a behavioral bias during uninformative trials, the presence of posterior alpha lateralization in an external reference frame when the touch was uninformative would have been suggestive of a role of alpha activity in tactile remapping proper. Its absence, in contrast, suggests that the lateralization observed in informative trials is involved in subsequent spatial processing.

Viewed together, externally coded, parietal alpha-band modulation occurs in a wide range of contexts, both during visual and tactile processing, prior and after stimulation. Rather than indexing, or even mediating, tactile-spatial transformation processes, it appears, therefore, to reflect a supramodal process related to orienting in space.

Tactile stimulation affected not only the power of alpha-band activity, but also its inter-regional coupling. Notably, we did not observe direct inter-hemispheric coupling of somatosensory and parietal cortex. We had hypothesized that information may be routed directly from somatosensory cortex of one hemisphere to the parietal cortex of the other when the hands are crossed. Such direct coupling could have been interpreted as a means of remapping somatotopic into external information through ad-hoc connectivity of the relevant parts of two differently coded spatial maps. Contrary to this hypothesis, during crossed-hand informative trials, connectivity manifested intra-hemispherically, contralateral to the external stimulus location and, thus, ipsilateral to the stimulated hand’s body side. Viewed together with the modulation of alpha-band activity in ipsilateral central electrodes by hand posture, and previous evidence indicating that primary and secondary somatosensory areas process both contra-an ipsilateral stimuli (Tamè et al., 2019), this result suggests that the somatosensory cortex ipsilateral to the anatomical side is involved in the transfer of information during postures that change regular, relative limb position.

### Behavior modulation precedes alpha-band modulation

Changes in oscillatory processes occurring while expecting, or after, a tactile stimulus have been shown to evolve around 400-1000 ms after the cue or stimulus (Jones et al., 2010; van Ede et al., 2011; Buchholz et al., 2013; Schubert et al., 2015, 2018). In the present study, central alpha-suppression associated with somatosensory processing was detected already 64 ms after stimulation, indicating that changes in alpha activity can occur at short latency. Critically, however, spatially-specific posterior alpha-lateralization occurred only after 150-300 ms and, thus, disassociated from the fast oculomotor search responses, especially when considering that saccade programming presumably finishes 100 ms prior to the overt saccade. If alpha-band lateralization were causal for saccade direction, it should precede, rather than follow, saccades. We explicitly modeled changes in alpha-band activity in relation to the subsequent saccadic behavior and did not observe modulation prior to saccade execution, suggesting against it being directly linked to the observed oculomotor behavior. Finally, modulatory effects of alpha-band connectivity first occurred more than 400 ms following tactile stimulation, which, just like power modulation, was later in time than the externally directed behavior. The consistent divergence of alpha-related modulation and externally oriented behavior suggests that the spatial processes mediated by posterior alpha-band lateralization are causally related neither tactile remapping, nor to exogenously oriented, fast overt behavior.

## Acknowledgments

We thank Christopher Lau and Lara Wurr for help with data acquisition. This work was supported by the German Research Foundation (DFG) through the Research Collaborative SFB936, project B1. TH was supported by a DFG Emmy Noether grant (He 8368/1-1).

## Author contributions

JPO, PK, and TH designed the experiment. JPO was responsible for the technical setup and acquired the data. JPO analyzed the data. JPO, PK, and TH wrote the manuscript. TH supervised the study.

